# Nanoparticle encapsulation enhances spatial distribution of Panobinostat to treat metastatic medulloblastoma via the intrathecal route

**DOI:** 10.64898/2026.03.31.715392

**Authors:** Oluwatobi Babayemi, Jon D. Larson, Sauradip Chaudhuri, Fred Valesquez, Janelle Morton, Chung-Fan Kuo, Lindsey K. Sablatura, Gerard Baquer, Michael S. Reagan, Sylwia Stopka, David I. Sandberg, Nathalie R. Agar, Eva Sevick-Muraca, Robert J. Wechsler-Reya, Rachael W. Sirianni

## Abstract

Medulloblastoma (MB) is an aggressive central nervous system (CNS) malignancy that primarily affects children and frequently exhibits metastasis to the leptomeninges of the brain and spinal cord. We developed a β-Cyclodextrin-poly(β-Amino Ester) nanoparticle system to deliver the histone deactylase inhibitor (HDACi) Panobinostat to MB by the intrathecal route. Various imaging methods were utilized to study nanoparticle and payload fate following infusion into the cerebrospinal fluid (CSF) of mice via cisterna magna or lumbar access points. Nanoparticles dramatically improved penetration of hydrophobic small molecules into distal regions of the spinal cord. Panobinostat-loaded nanoparticles were effective at treating patient-derived MB, activating pharmacodynamic targets, slowing growth of the primary tumor, decreasing incidence of metastasis at the time of death, and ultimately prolonging survival. These studies provide insight into the mechanisms mediating transport of colloids and therapeutic molecules in the subarachnoid space and highlight new approaches for treating metastatic disease in the CNS.

## Introduction

Medulloblastoma (MB) is a rare but highly aggressive central nervous system (CNS) malignancy that predominately affects children and adolescents.^1^ Standard of care for MB includes surgical resection, chemotherapy, and high-dose cranio-spinal radiation. Many children will survive their disease, however, long-term outcomes remain poor due to treatment-associated damage to the developing CNS.^2^ This has been a particular challenge for Group 3 and Group 4 MB, which have a propensity to metastasize to the leptomeningeal membranes surrounding the brain and spinal cord. Leptomeningeal metastasis (LM) remains difficult to treat by conventional approaches and is considered untreatable in a recurrent setting, which highlights the urgent need for better therapies.^3^

Here, we focused on therapeutic approaches for the delivery of a histone deacetylase inhibitor (HDACi), Panobinostat.^4,5^ Panobinostat is a lipophilic, poorly water-soluble small molecule that has received substantial attention in recent years for its possible use in the treatment of CNS malignancies, including glioblastoma (GBM) (NCT05324501, NCT00859222, NCT00848523), diffuse intrinsic pontine glioma (DIPG) (NCT04341311, NCT02717455, NCT03632317, NCT03566199, NCT05009992), and by us in pediatric MB (NCT04315064). Panobinostat exerts significant anticancer effects against each of these tumor types *in vitro*, however, delivery remains a challenge. We previously showed that although intravenously administered Panobinostat can cross the blood-brain barrier (BBB) and blood-spinal cord barrier (BSCB) to reach the CNS parenchyma in mice at detectable levels, its free fraction in brain tissue is both regionally variable and below the optimal therapeutic ranges that are expected based on *in vitro* potency of this compound.^6,7^ Poor bioavailability, enzymatic degradation, protein binding, and limited tissue penetration are impediments to the translation of Panobinostat into an effective clinical therapy.^6,7^

The delivery challenges associated with Panobinostat use are common to other drugs within the HDACi family, and nanoparticle approaches for enhancing drug delivery may offer one solution.^8–11^ Nanoparticles provide several advantages for improving delivery of small molecules to the CNS, including protection of active agents from physiological environments and sustained release, which enhances total tissue exposure to drug. We previously developed a generalizable strategy for the encapsulation and slow release of HDACi from polymeric nanoparticles composed of poly(beta-amino ester)-beta-cyclodextrin (CDN).^11^ In this prior work, we showed that encapsulation of Panobinostat within the CDN system (i.e., pCDN) enabled a 30-fold higher deliverable dose when administered via convection-enhanced delivery into brains of mice bearing orthotopic GL261 tumors. The improvement in delivered dose resulted in a higher intratumoral Panobinostat concentration compared to when mice received Panobinostat solubilized within beta-cyclodextrin (an approach that does not offer controlled release), which ultimately yielded treatment efficacy.^11^

Here, we sought to further develop the pCDN system for treatment of MB by focusing on intrathecal administration, i.e., direct infusion of nanoparticles into the cerebrospinal fluid (CSF) that fills the subarachnoid spaces of the brain and spinal cord.^12^ First, the CDN network was engineered to ensure nanoparticle stability in CSF. Next, we evaluated the spatial distribution of hydrophobic molecules within CNS tissues following administration of nanoparticles into the cisterna magna of healthy mice. Finally, we studied the therapeutic efficacy of pCDNs in mice bearing an orthotopic, patient-derived xenograft (PDX) model of Group 3 MB exhibiting LM. We hypothesized that nanoparticle encapsulation of Panobinostat would enhance delivery to the CNS via the intrathecal route, which we predicted would enable treatment of patient-derived MB exhibiting LM. Our results show that nanoparticle encapsulation dramatically improves both drug tolerability and distribution of encapsulated agents within the CNS and that the growth of tumors is slowed following intrathecal administration of nanoparticles, leading to improved survival compared to mice who received drug-empty control nanoparticles. Taken in sum, this work presents a newly engineered, stabilized version of CDN nanoparticles for HDACi delivery to the CNS and also advances our mechanistic understanding of how to effectively engineer colloidal systems for intrathecal drug delivery.

## 2. Results

### 2.1 Synthesis and Characterization of CDN-5 Nanoparticles

The cyclodextrin network (CDN) polymer consists of beta cyclodextrin covalently linked to a poly(beta amino ester) (PBAE) backbone and functionalized with poly(ethylene glycol) (PEG).^11^ By varying crosslinker type, length, and concentration, we generated a variety of amphiphilic structures that self-assemble into a nanoparticle following addition of a hydrophobic payload (**Figure 1**). The CDN-5 structure is similar to CDN-4, which we reported previously,^11^ excepting the following alterations: the molecular weight of the PEG was shortened from 550 to 221 Da, the concentration of crosslinker was reduced by half, and side group functionalities were modified to possess a –COOH moiety. Together, these modifications yielded a more compact nanoparticle core with enhanced hydrophilicity on the surface. CDN-5 was additionally surface-modified through Michael addition with NODAGA, permitting chelation of ^64^Cu for subsequent imaging purposes. We note here the nomenclature for CDN polymer following its formation into nanoparticles: *surface modification-payload-CDN_version#_*.

**Figure 1.**
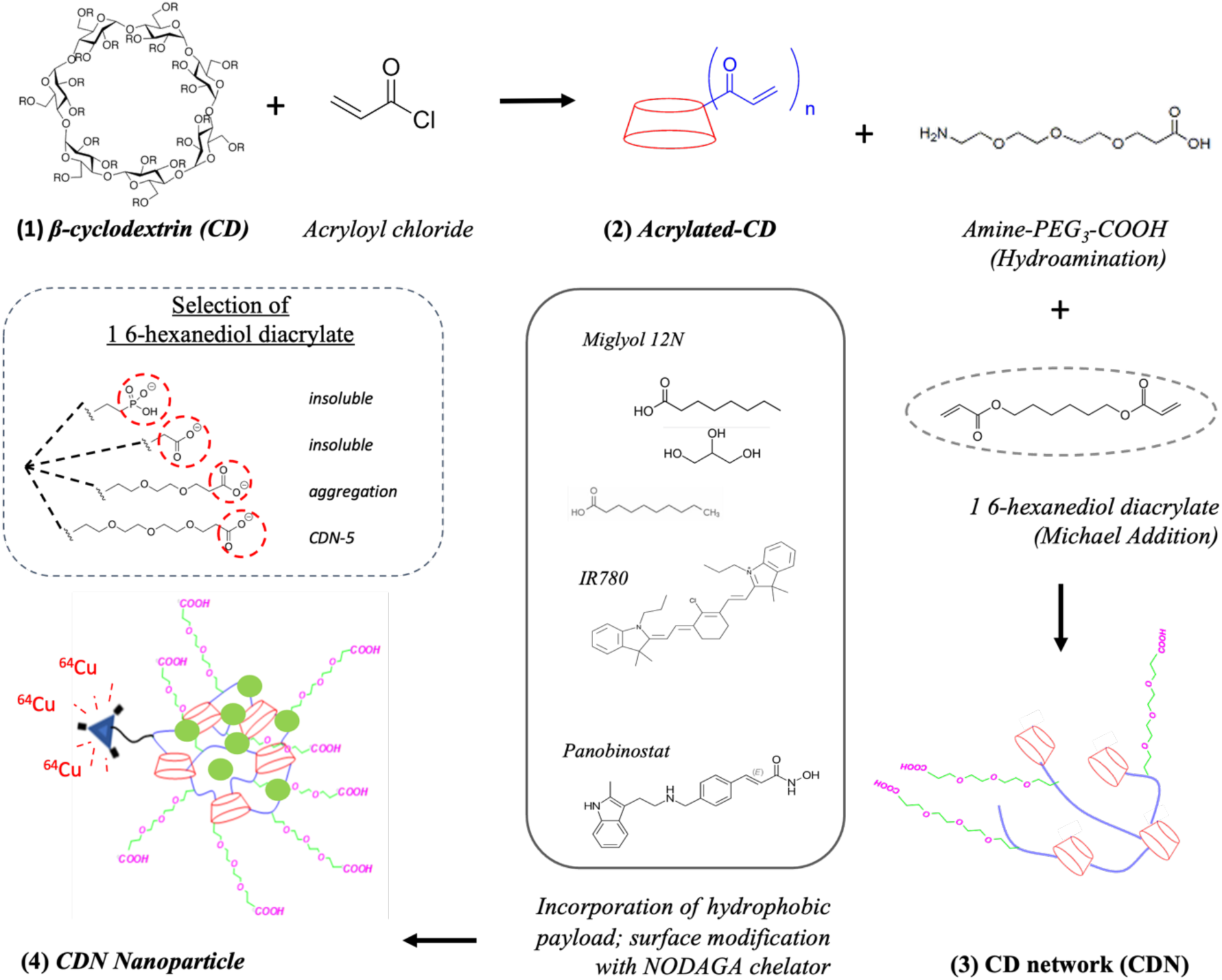
**CDN-5 was developed through rational design**. CDNs consist of a poly(beta-amino ester) (PBAE) backbone crosslinked to beta-cyclodextrin (CD) and polyethylene glycol (PEG). A general formulation scheme follows. The addition of acryloyl chloride to **(1) β-cyclodextrin** yields **(2) acrylated CD**. Michael addition of acrylated-CD with PEG and ester constituents yields a water soluble, cross-linked **(3) cyclodextrin network (CDN).** The more stable CDN network (i.e., CDN-5) was engineered by modulating hydrophobic interactions in the particle core via adjustment of length and monomer end groups. Formulation proceeds via self-assembly, by doping the constituent CDN with hydrophobic payload in water with a small quantity of DMSO; the inclusion of a hydrophobic small molecule drives formation of the **(4) CDN nanoparticle**, which was surface functionalized with NODAGA to enables chelation of Cu-64 for multifunctional delivery and imaging.

The synthesis of these materials with expected chemical functionalities was confirmed with NMR analyses (see **SI1**). Following addition of hydrophobic payload, solutions shifted from turbid to translucent, and CDN polymers were observed to self-assemble into discrete, drug loaded nanoparticles size, surface charge, and drug loading suitable for *in vivo* administration (**Figure 2A**). Solubilization of Panobinostat in beta-cyclodextrin (i.e., pCD, which is similar to the clinically used formulation^13^) yielded very poor loading on a mass basis (0.38 ± 0.13 w/w%). Similar to our prior report, Panobinostat loaded CDN-4 nanoparticles (i.e., pCDN_4_) were highly loaded (27.2 ± 5.0 w/w%) and possessed a favorable hydrodynamic diameter (283.4 ± 51.8nm) and zeta potential (1.2 ±1.7 mV) for *in vivo* use, while Panobinostat loaded CDN-3 nanoparticles (i.e., pCDN_3_) possessed moderately lower loading (17.5 ± 1.3 w/w%), much smaller diameter (75.2 ± 9.4nm) and a more positive zeta potential (20.3 ± 1.9 mV). Panobinostat loaded CDN-5 nanoparticles (i.e., pCDN_5_) yielded similar loading (19.0 ± 4.3 w/w%), slightly larger diameter (337.0 ± 48.3), and a lower but still close to neutral zeta potential (-7.9 ± 2.0 mV). Following incorporation of IR780 and modification of CDN-5 with NODAGA for chelation of ^64^Cu (i.e., irCDN_5_), nanoparticles retained these favorable properties, with a similar hydrodynamic diameter and zeta potential to other formulations (353.7 ± 12.0 nm and -10.9 ± 2.7 mV, respectively). Transmission electron microscopy (TEM) analyses demonstrated that nanoparticles fabricated from CDN-5 are non-aggregated and possess a smooth surface morphology (**Figure 2B**). These data affirm the ability of the CDN polymer to self-assemble into nanoparticles following addition of a hydrophobic payload. Furthermore, they show that our modifications, which confer multi-modal imaging functionality, do not significantly change the colloidal features of resultant nanoparticles.

**Figure 2.**
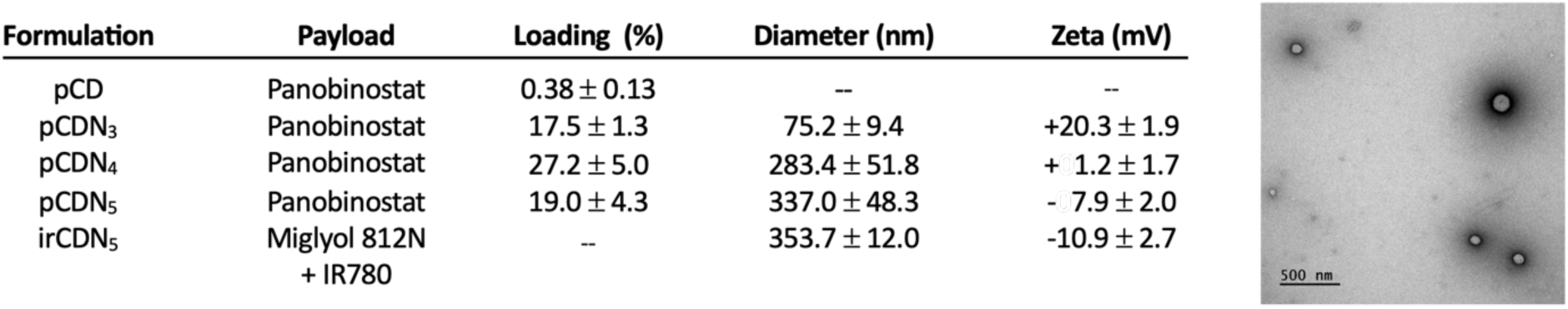
CDN nanoparticles form via self-assembly following the addition of hydrophobic payloads. Various formulations were developed in the course of this work, loading example molecules of Panobinostat, IR780, and Miglyol® 812 into CD or the CDN-3, CDN-4, or CDN-5 networks. (A) Physical characterization of the delivery systems presented in this work. Error represents the standard deviation of 3 experimental replicates. (B) Transmission Electron Microscopy (TEM) images of pCDN_5_ show that nanoparticle morphology is spherical with smooth surfaces. Scale bar = 500nm.

To assess the colloidal stability of pCD, pCDN_3_, pCDN_4_, and pCDN_5_, formulations were freshly prepared, suspended in aCSF of varying pH, and subjected to DLS analysis at regular intervals for up to 12 days. Here, “instability” refers to an increase in hydrodynamic diameter above 1μm. These experiments yielded several important observations (**Figure 3**). First, the pCD formulation was particularly unpredictable compared to the nanoparticle alternatives, with a large degree of aggregation observed in all tested conditions, including immediately after suspension in high pH aCSF. pCDN_3_ was stable in both acidic and neutral pH, whereas time-dependent aggregation was observed in alkaline pH. pCDN_4_ was unstable in both acidic and neutral pH, and time-dependent aggregation was observed in alkaline pH. In contrast, pCDN_5_ maintained a stable hydrodynamic diameter in both neutral and alkaline pH. Some aggregation of pCDN_5_ was observed at day 6 in acidic conditions only. The eventual aggregation at later time points is likely due to the release of Panobinostat into the incubation media, which will result in disassembly of the organized nanoparticle into free polymer that then forms larger aggregates. pCDN_5_ stability in neutral media was considered excellent, with no evidence of aggregation even after 12 days of incubation at 37°C. These data affirm that the CDN-5 structure improves colloidal stability of drug loaded nanoparticles in aCSF relative to native cyclodextrin or CDN-4.

**Figure 3.**
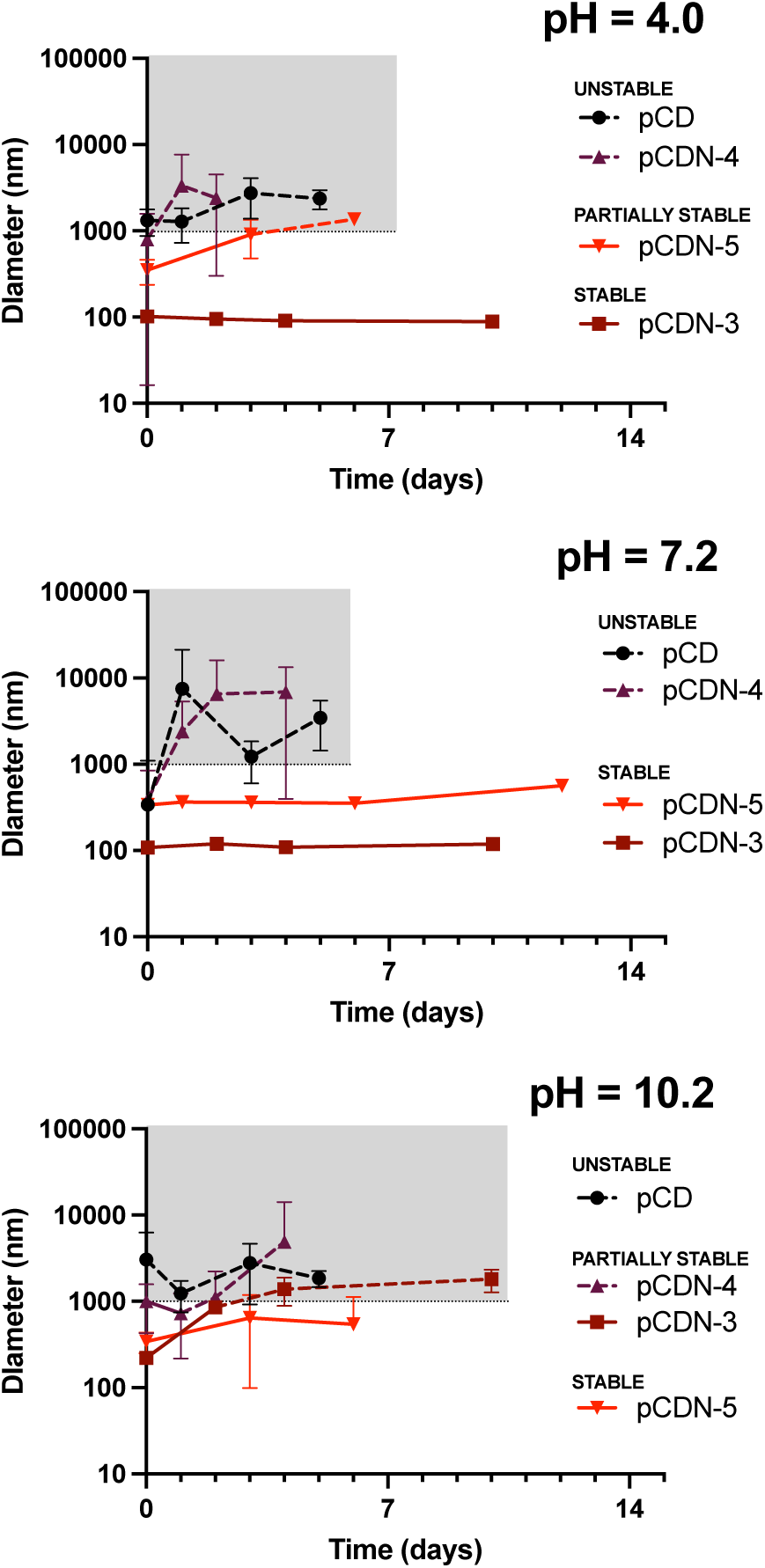
CDN_5_ nanoparticles demonstrate enhanced in vitro stability relative to alternative formulations. Drug loaded particles were freshly prepared in aCSF with pH adjusted to yield (A) acidic, (B) neutral, or (C) alkaline conditions. pCD and pCDN_4_ aggregated immediately, regardless of pH. pCDN_3_ was highly stable in both acidic and neutral pH aCSF, with moderate instability in alkaline pH over time. pCDN_5_ exhibited good stability in neutral and alkaline pH, with mild aggregation observed at later time points in acidic pH. Aggregation of pCDN_5_ in acidic media was observed only at the latest time point (day 6), at which point the system is expected to disassemble due to release of drug. Aggregation is defined as an aqueous diameter of >1000nm. Data describing samples for which aggregation was observed are represented by a dashed line. Data points describe mean with standard deviation for a minimum of 3 replicates.

### 2.2 Nanoparticle and Payload Fate in the Subarachnoid Space

To assess nanoparticle fate following *in vivo* administration, we developed methods to radiolabel CDN-5 with the gamma-emitting radioisotope ^64^Cu using NODAGA as a chelator, yielding ^64^Cu-irCDN_5_, a system that can be imaged with Positron Emission Tomography (PET). The optimized approaches yielded excellent ^64^Cu chelation efficiency (95.2±0.83%, n=4 independent batches) and reliably high radioactivity yields (1.1±0.24 µCi/µL, n=4 independent batches). Following administration into the CSF of healthy mice, ^64^Cu-labeled CDN_5_ that was not assembled into nanoparticle form (i.e., lacking hydrophobic payload) remained localized exclusively to the cisternal injection site. By contrast, ^64^Cu-irCDN_5_ nanoparticles, which are loaded with IR780 payload and therefore assembled, were observed to rapidly distribute throughout the subarachnoid space (**Figure 4**). These data provide empirical confirmation that the CDN_5_ nanoparticle system is stable *in vivo* and support the expectation that movement through the subarachnoid space is improved by assembly of the polymer into a stable nanoparticle form.

**Figure 4.**
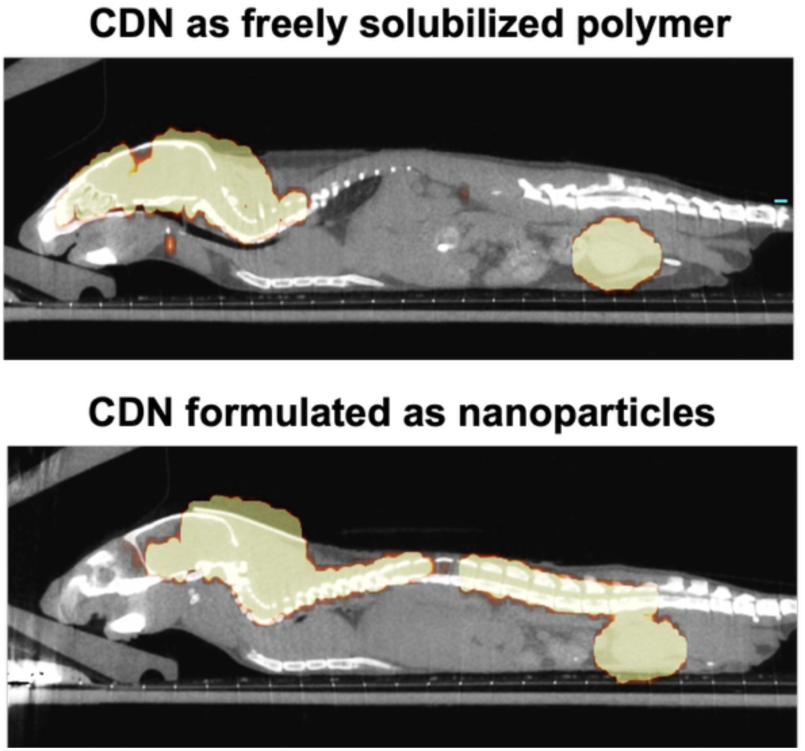
**Nanoparticles must be assembled for efficient transport within the subarachnoid space**. When CDN-5 was administered via the IT-CM route as a freely soluble network, delivery was confined to the brain subarachnoid space immediately adjacent to the cisterna magna. In contrast, once the CDN-5 was assembled into nanoparticle form through addition of the hydrophobic payload, distribution was reliably observed throughout the subarachnoid space of both the brain and spinal cord.

Intrathecal (IT) delivery is defined as administration of a substance directly into CSF, and this can be achieved through multiple sites, including ventricular, lumbar (IT-L), and cisterna magna (IT-CM) access points. To identify the optimal route for intrathecal administration, we evaluated the spatial distribution of nanoparticles following injection via the IT-L vs IT-CM routes with PET/CT imaging. These experiments yielded several important observations. For the IT-L group, particles were observed primarily within the L5/L6 region. Some nanoparticles transited rostrally to reach the brain subarachnoid space, although the relative concentration of nanoparticles within the brain subarachnoid space in the IT-L group was much lower than in the IT-CM group (**Figure 5A**). For the IT-CM group, nanoparticles were initially observed at a high concentration near the base of the brain and cervical spinal cord, after which they distributed through the ventral aspects of the brain subarachnoid space and throughout the spinal cord subarachnoid space. In some subjects, coverage of the brain subarachnoid space was complete, encompassing both dorsal and ventral aspects of the brain, while other subjects exhibited primarily ventral distribution, and no subject exhibited primarily dorsal distribution. Quantification of these data (**Figure 5B**) yielded major differences in the patterns of delivery that are achieved by different access points. We observed a characteristic pattern of distribution for IT-CM administered nanoparticles in which high signal was observed in the upper thoracic and cervical regions of the spinal cord, while a lower signal was observed in the lumbar spinal cord; a secondary peak in nanoparticle delivery via the IT-CM route is observed at the base of the spine, near the cauda equina. The reduction in signal observed in the lumbar region of the spinal cord was observed regardless of subject, experimenter, and experimental days (not shown). In contrast to IT-CM, IT-L administration yielded a spatially consistent, low level delivery of nanoparticles to thoracic, lumbar, and sacral regions of the spinal cord, with inconsistent and incomplete delivery to the brain subarachnoid space (**Figure 5B**). Quantifying the fraction delivered to the brain out of total CNS delivery highlights major differences between these routes of administration: the brain ROI contains ∼74-76% of the total delivered dose for the IT-CM route, whereas the brain contains ∼14-16% of the total delivered dose for the IT-L route, whether assessed at early or late time points (**Figure 5C**). IT-CM therefore offers advantages over IT-L for brain delivery, and we focused our work on IT-CM moving forward.

**Figure 5.**
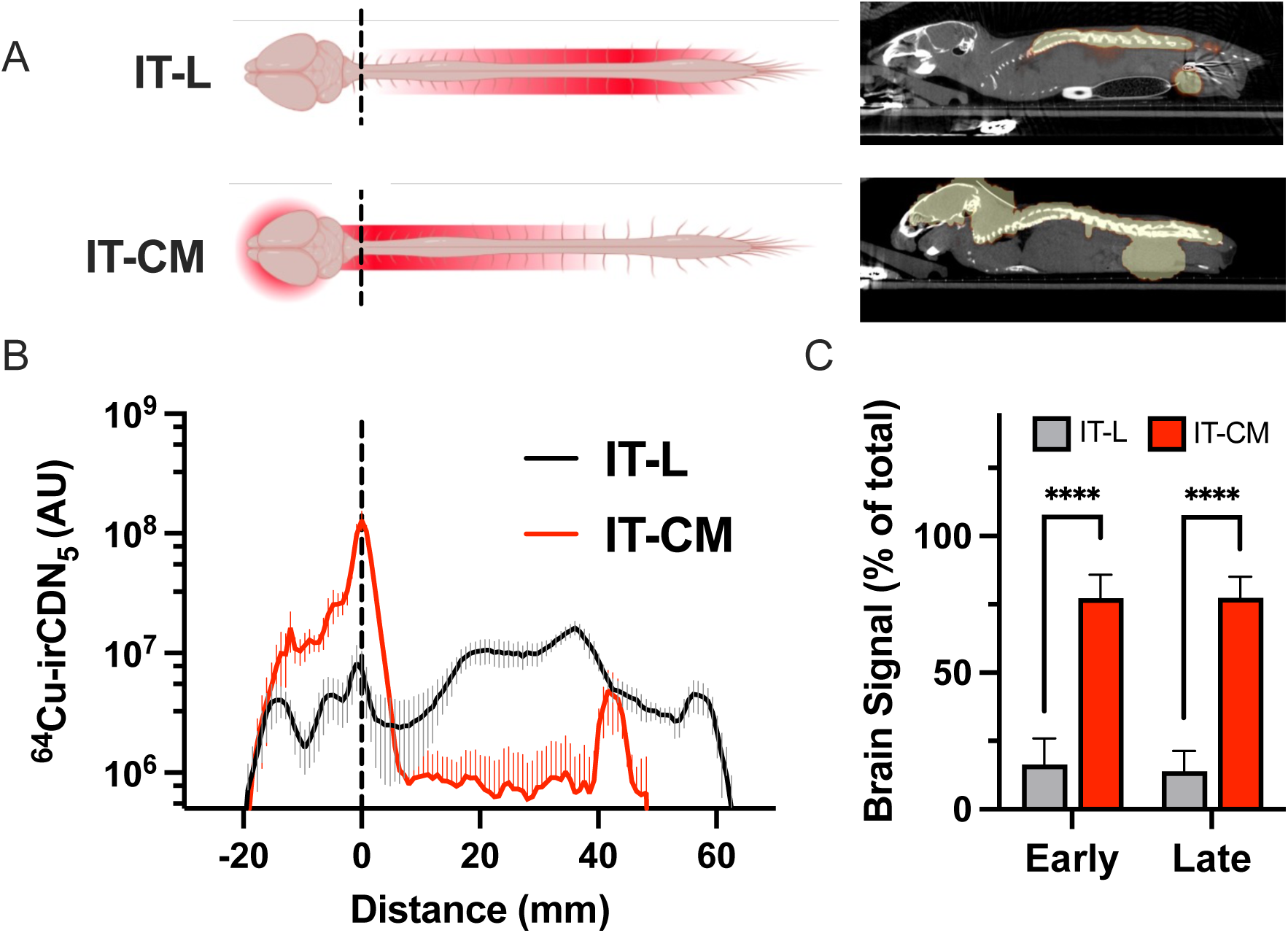
**IT-CM is a preferred route of administration over IT-L for achieving delivery to the brain subarachnoid space**. (A) When ^64Cu^CDN_5_ nanoparticles were administered intrathecally via the lumbar route (IT-L), PET imaging demonstrated that delivery remained concentrated near the injection site, and nanoparticles achieved minimal access to the subarachnoid space of the brain. In contrast to the IT-L route, administration of nanoparticles intrathecally via the cisterna magna (IT-CM) yielded widespread distribution within the subarachnoid space in both the brain and the spinal cord. (B) Distribution of nanoparticle signal along the neuroaxis was quantified from the imaging data (n=4 per group). Data were aligned to the IT-CM injection point. We note that while the olfactory bulb to cisterna magna distances are fixed, the apparent length of the spinal cord varies depending on how the mouse is positioned (flat versus curved back), and so distributions from the cisterna magna to cauda equina exhibit variability in the apparent termination of signal at the end of the spinal cord. (C) The relative fraction of nanoparticles delivered to the brain vs other sites was quantified across experiments by aggregating early (<40 min) vs late (>40 min) time points. Values represent the mean with error bars representing standard deviation.

The ability of CDNs to deliver hydrophobic payloads to the CNS was first evaluated by studying the *ex vivo* fluorescence of IR780 following IT-CM administration in free form or from ^64^Cu-irCDN_5_ nanoparticles. When IR780 was administered in free form, signal was typically detected adjacent to the injection site with minimal signal at distal locations, save for the base of the spinal cord at the cauda equina (**Figure 6a**). Total detectable signal was relatively low and could not be visualized in some subjects. In contrast, when IR780 was delivered from nanoparticles, signal was much brighter and widespread throughout the neuroaxis (**Figure 6A**). These patterns of delivery are also evident on visual assessment of fluorescence within the spinal cord (**Figure 6B**). Although a gap in nanoparticle distribution was observed in the lumbar region of the spinal cord (**Figure 5B**), payload distribution was relatively uniform across this same region (**Figure 6A**). Calculation of the spinal cord AUC further highlights the advantages of nanoparticle delivery; the AUC for freely administered IR780 was 690, 4700±310, and 6800±440 AU*mm at ∼7, 37, and 67 minutes, respectively, compared to the AUC for irCDN_5_ of 23,000±450, 26,000±700, and 39,000±450 AU*mm at ∼52, 82, and 112 minutes, respectively. Ultimately, both the total delivery and the spatial distribution of IR780 were markedly higher when delivered from nanoparticles compared to administration in free form.

**Figure 6.**
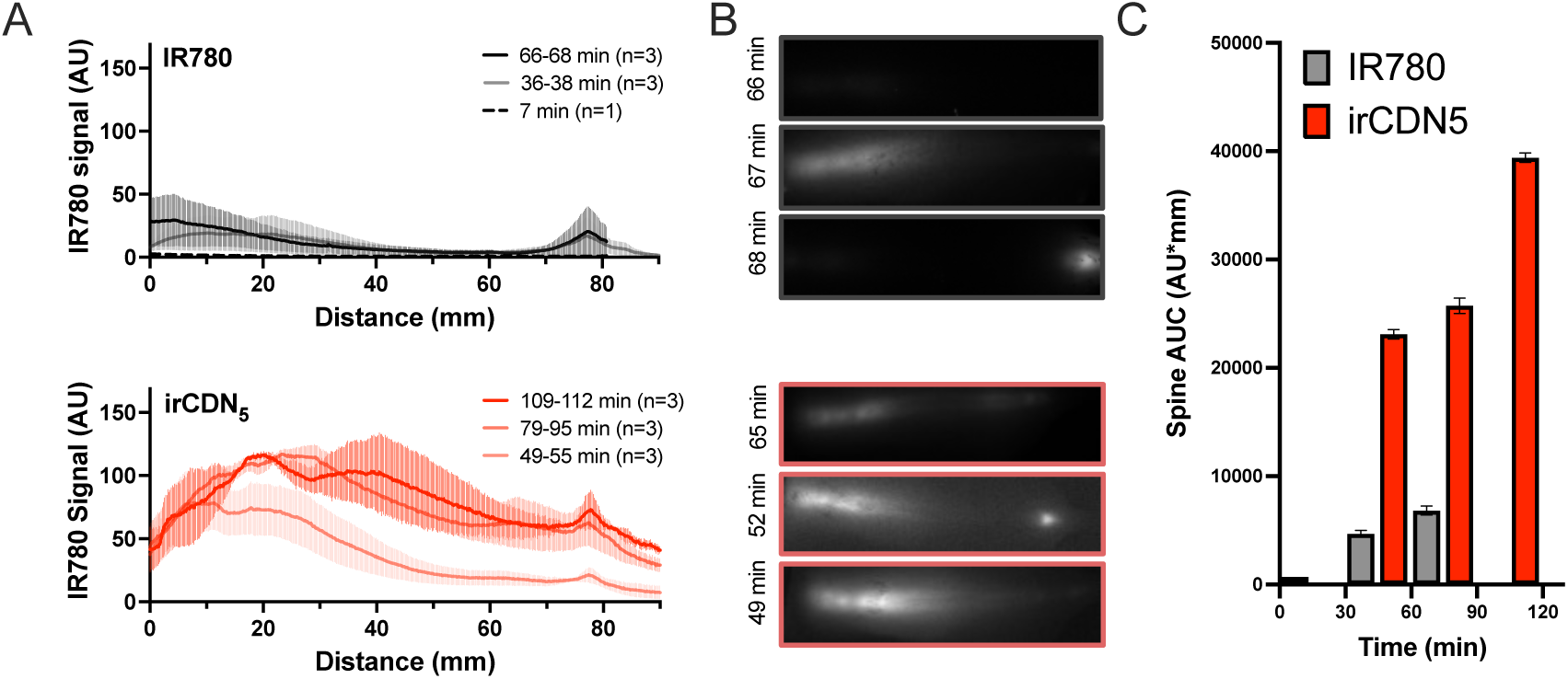
**The spatial distribution of IR780 along the neuroaxis is markedly enhanced by encapsulation within a CDN-5 nanoparticle**. Mice received IT-CM injections of either freely solubilized IR780 or irCDN_5_. At various time points after substance administration, mice were imaged with a near-infrared fluorescent camera system. (A) The level of IR780 was quantified as a function of rostral-caudal distance along the neuraxis, with a distance of 0 corresponding to the upper cervical region of the spinal cord. (B) Representative examples of IR780 fluorescence in whole spinal cord. (C) The total signal within the spine, i.e., AUC, was calculated from the data presented in subpanel (A). Group sizes and exact imaging times are provided in the figure caption of subpanel (A).

### 2.3 Treatment studies

Panobinostat is a model HDACi that faces significant delivery challenges in the body. As a poorly water-soluble and lipophilic agent, Panobinostat cannot be administered to CSF in free form. These experiments therefore focused on comparing the drug distributions resulting from administration of pCD—a solubilization approach that is used clinically but does not offer controlled release, and that our data show is relatively unstable in aCSF— versus administration of pCDN_5_ — which our data show is stable in aCSF. We predicted that encapsulation within pCDN_5_ would facilitate transport of Panobinostat payload to distal regions of the CNS.

Tolerability was first evaluated in non-tumor bearing C57BL/6 mice (**Figure SI3**). We did not reach an MTD via the intravenous route; intravenous tolerability instead was limited by maximum feasible dosing, which was 80μg for pCD and 2000μg for pCDN. In contrast, via the IT-CM route, the single-dose MTD for pCDs was 2μg, compared to 8μg for pCDN_5_, emphasizing the ability of controlled release systems to decrease drug-associated toxicity. These differences were heightened in a multidose setting, where the MTD for twice-weekly administration by the IT-CM route was determined to be 0.5μg for pCD and 6-7μg for pCDN_5_. Thus, nanoparticle encapsulation improved tolerability of Panobinostat by 4-fold in a single-dose setting and by 12-14-fold in a multidose setting.

The spatial distribution of Panobinostat was evaluated as a function its formulation using MALDI mass spectrometry imaging (MALDI-MSI). Either pCDs or pCDN_5_ nanoparticles were administered by the IT-CM route at their individual acute-dose MTD (2 and 8μg, respectively) to healthy C57BL/6 mice. The brains and spinal cords of these mice were extracted 2 hours later for MALDI MSI analyses, which yielded spatial maps of drug concentration in tissue (**Figure 7A**). At this time point, the total concentration of Panobinostat detected in whole-brain scaled linearly with administered dose, yielding 1.5±1.4 µM and 6.3±7.1 µM for pCD versus pCDN groups (**Figure 7B**). Similarly scaled enhancements were observed for a periventricular ROI, yielding 15±10 µM and 60±72 µM for pCD vs pCDN groups. Gross patterns of spatial distribution were similar to what we observed for IR780: the highest concentration of Panobinostat was detected immediately adjacent to the injection site, with additional delivery concentrated along the ventral aspect of the cerebellum and throughout parenchymal tissues adjacent to the 4^th^ ventricle. Some evidence for increased penetration of Panobinostat into ventricular-adjacent tissue was observed for the pCDN group, with an increase in the effective volume of distribution from 6.1±1.1 to 9.1±1.9% (i.e., an ∼50% increase in volume of distribution), although these differences were difficult to quantify due to the complex, small tissue geometry of the ventricles and were not statistically significant.

**Figure 7.**
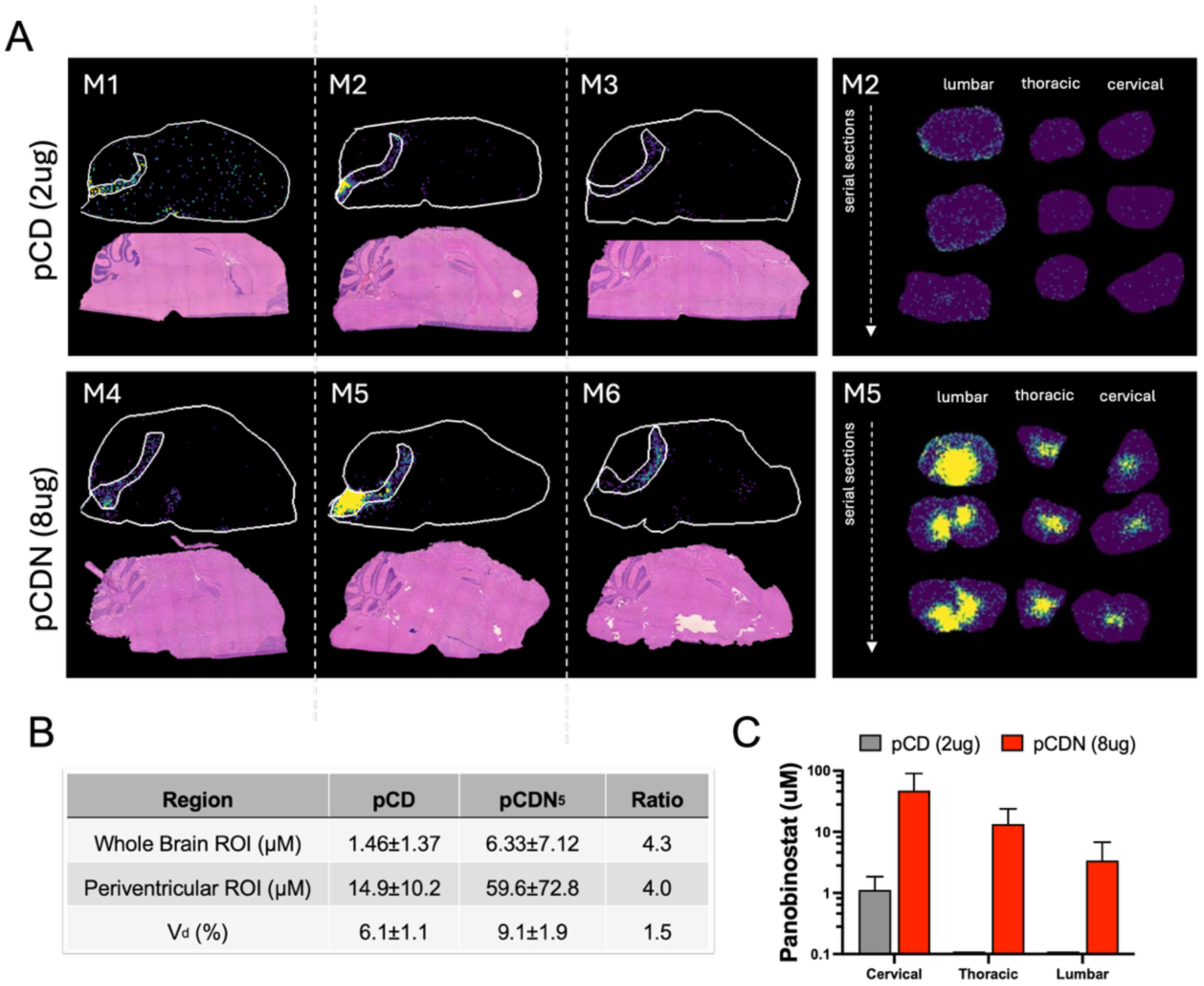
The spatial distribution of Panobinostat is markedly enhanced by encapsulation within a CDN-5 nanoparticle. Mice received IT-CM injections of either pCD or pCDN_5_. Two hours after substance administration, tissues were extracted for MALDI-MSI analyses. (A) Paired drug distribution and H&E sections show drug distribution in the brain (left) and spinal cord (right). Spinal cord sections show M2 and M5, which were the subjects with the highest brain signal of Panobinostat for pCD and pCDN5 treatment, respectively. Whole Brain and Periventricular ROIs are outlined in white. (B) Panobinostat levels were quantified in each region of interest, and the ratio between the treatment groups as well as the apparent volume of distribution was calculated. (C) Panobinostat levels in the spinal cord were calculated for cervical, thoracic, and lumbar regions. Experiments represent tissues obtained from a total of 6 mice, with data presented as mean plus/minus standard deviation.

Examination of drug distribution distal to the injection site revealed dramatic differences in Panobinostat penetration into the spinal cord, depending on formulation. For the pCD group, Panobinostat was only detectable in one of three subjects within the cervical region of the spinal cord; no drug could be detected in distal locations. In contrast, for the pCDN_5_ group, Panobinostat was detected at very high concentrations throughout the spinal cord, achieving a concentration of 47.2µM in cervical spinal cord and declining in an exponential fashion to reach 3.4µM in the lumbar region. Interestingly, the highest drug concentrations in the spinal cord were observed in the medial aspect, near the central canal, rather than in the lateral aspects that are immediately adjacent to the leptomeninges (**Figure 7A**). We previously reported nanoparticle transport through the central canal following IT-CM administration,^8^ and the data presented here suggest that flow of nanoparticles through the central canal could facilitate drug delivery to the parenchyma of the spinal cord. Additional MALDI-MSI data, including in tumor bearing mice, are provided in *Supplementary Data* (SI Figure 2); these additional data are more variable but consistent with the observations reported here, with higher levels of drug detected in CNS tissues, including tumor ROI, for mice that received pCDN5 compared to mice that received pCD.

The next series of experiments tested whether IT-CM administered pCDN_5_ would activate the expected pharmacodynamic targets. H3K9ac and FOXO1 expression levels are known biomarkers that are responsive to Panobinostat exposure.^14^ Mice harboring primary Med-411FH tumors in the cerebellum and exhibiting metastasis were treated with either 8μg of pCDN_5_ nanoparticles or an equivalent mass dose of drug-empty CDN_5_ (bCDN_5_) delivered by the IT-CM route. Treatments occurred at an advanced, i.e., moribund, stage of tumor growth, and cerebellum and spinal cord tumor cells were harvested 6 hours later to examine H3K9ac protein and *FOXO1* mRNA expression. We observed a significant increase in H3K9ac protein and *FOXO1* mRNA expression in pCDN_5_-treated cerebellum tumor samples relative to bCDN_5_ controls (**Figure 8A-C**). pCDN_5_-treated spinal cord tumor samples showed a modest increase in *FOXO1* mRNA expression that was not statistically significant (**Figure 8D**).

**Figure 8.**
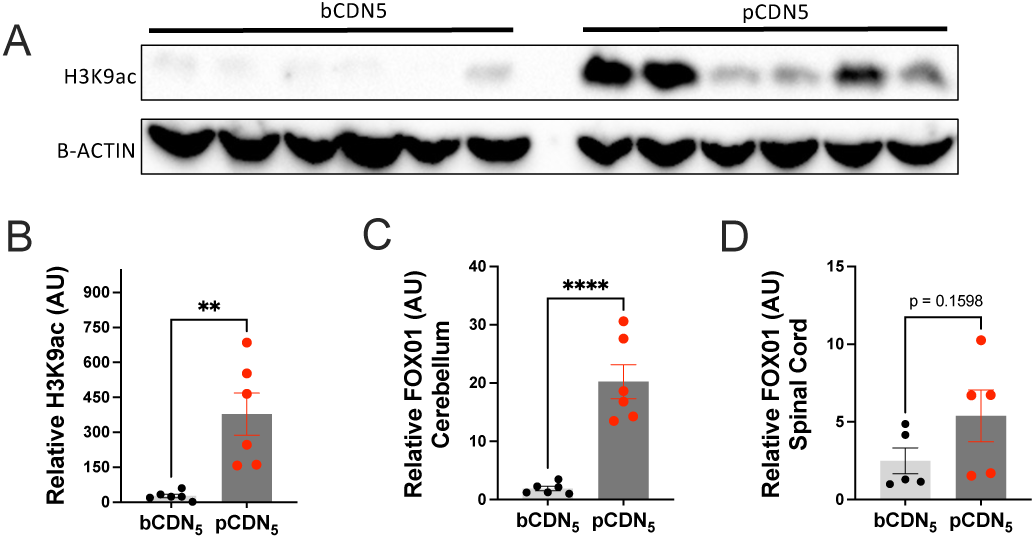
IT-CM administration of pCDN_5_ to mice harboring metastatic MB xenografts activates pharmacodynamic targets *in vivo*. Mice bearing late-stage Med-411FH tumors exhibiting LM were treated with empty (non-drug loaded) bCDN_5_ or Panobinostat-loaded pCDN_5_ nanoparticles. Tumors were harvested and analyzed 6 hours after treatment. (A) Western blot showing H3K9ac protein levels in cerebellum tumor cells treated with bCDN_5_ or pCDN_5_ nanoparticles. (B) Quantification of relative H3K9ac normalized to B-ACTIN from subpanel (A). (C, D) qRT-PCR showing *FOXO1* mRNA quantification relative to *B-ACTIN* in tumor cells extracted from the cerebellum (C) or spine metastasis (D). Data in (B-D) represent the mean value of 6 mice per group, with error bars representing the standard error of the mean. P-values are reported on the following scale: *<0.05, **<0.01, ***<0.001, ****<0.0001.

Given the positive pharmacodynamic response of tumors to a single dose of pCDN_5_, we next examined whether repeat pCDN_5_ treatment could exert a therapeutic effect on orthotopic Med-411FH tumors when supplied at the multi-dose MTD. As before, tumors were induced by transplanting cells into the cerebellum, which is a model that reliably yields metastatic lesions. Either 6μg of pCDN_5_ or an equivalent bCDN_5_ mass were administered by the IT-CM route on a twice-weekly basis (**Figure 9A**). We observed that treatment with pCDN_5_ resulted in a significant reduction in tumor doubling time in both brain and spinal cord regions of interest (**Figure 9B**), which ultimately yielded an increase in median survival for pCDN_5_ treatment compared to control (p = 0.0039, Mantel-Cox test) (**Figure 9C**). In each week of treatment, the surviving mice received bioluminescent imaging to evaluate tumor growth and extent of metastasis. Mice were considered to have metastatic disease when the bioluminescent signal exceeded XXX Mice that received bCDN_5_ tended to have a higher occurrence of LM that could be identified by bioluminescence imaging (**Figure 9D**). For example, when assessed at week 6 of treatment, 43% of bCDN_5_ treated mice exhibited or had previously exhibited (prior to endpoint) LM compared to 16% of pCDN_5_ treated mice; during week 7, these numbers rose to 92% and 56% of bCDN_5_ and pCDN_5_ treated mice, respectively. These data establish that intrathecally administered pCDN_5_ is efficacious for treatment of orthotopic, patient-derived MB exhibiting LM.

**Figure 9.**
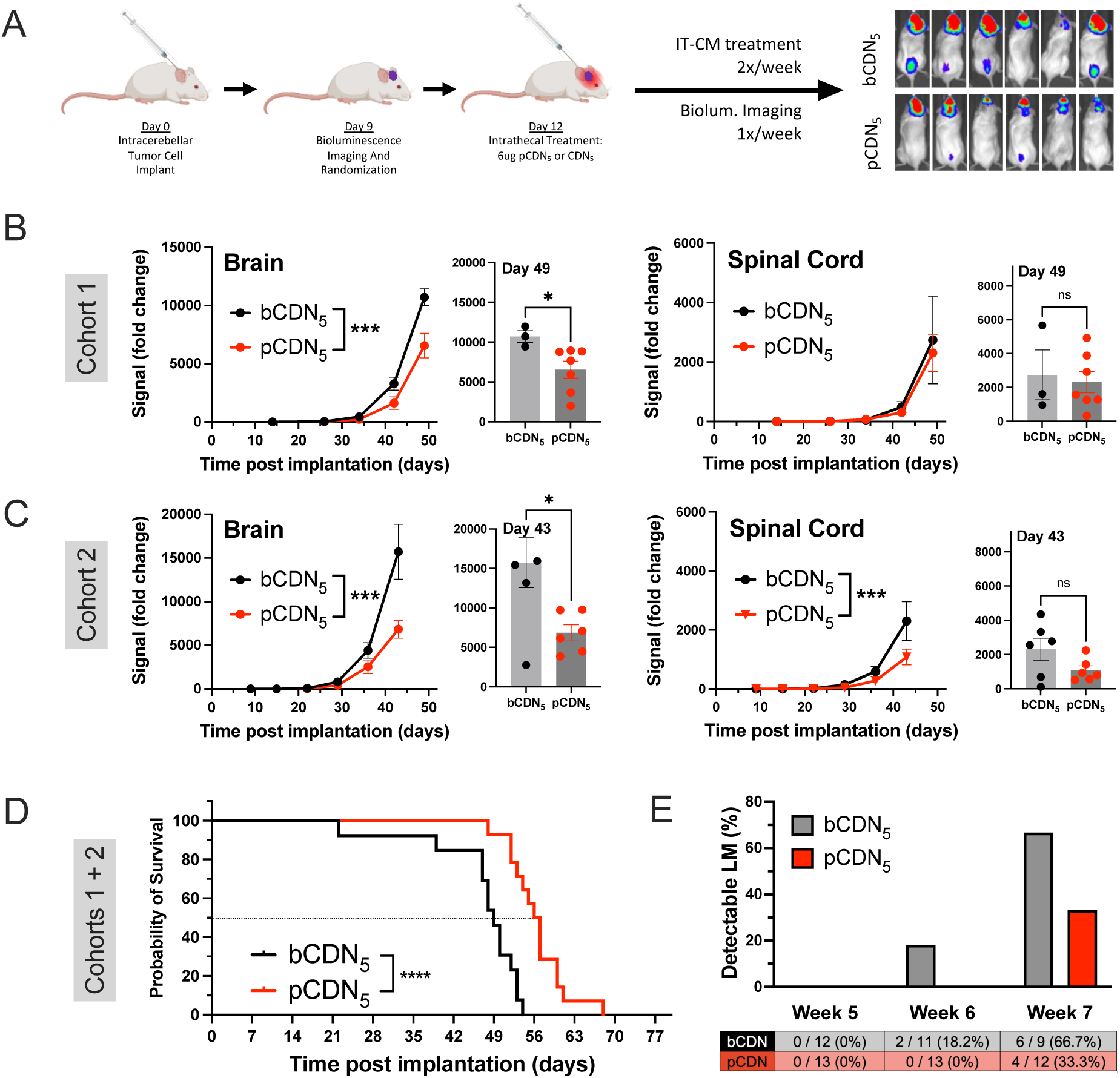
IT-CM-mediated pCDN_5_ treatment slows tumor growth and prolongs survival of mice harboring patient-derived metastatic MB xenografts. Med-411FH PDX tumor-bearing mice treated by IT-CM administration of bCDN_5_ or pCDN_5_ (6ug). These studies were conducted in two cohorts, consisting of n = 7 and 8 (cohort 1) and n = 6 and 6 (cohort 2) for bCDN_5_ and pCDN_5_, respectively. (A) Strategy to evaluate pCDN_5_ treatment efficacy, including 2x/week treatments and 1x/week bioluminescent imaging. (B) Tumor growth was measured by the fold-change of bioluminescent signal relative to pre-treatment signal. These data are presented as independent cohorts due to minor variation in imaging day. (C) Kaplan-Meier survival analysis of both cohorts yields a significant prolongation of survival for mice treated with pCDN_5_. (D) The fraction of mice exhibiting LM based on bioluminescent assessment was determined for each week of the study. Data represent the mean plus/minus standard error for a minimum of 3 subjects. Statistical testing for the Kaplan-Meier plot was conducted by Log-rank (Mantel Cox) test. P-values are reported on the following scale: *<0.05, **<0.01, ***<0.001, ****<0.0001. Mouse images created with Biorender.com.

## 3. Discussion

Medulloblastoma is the most common malignant brain tumor in children, and long-term outcomes remain poor, in part due to limited treatment options.^15^ LM typically cannot be surgically resected, and high doses of radiation needed to treat or prevent metastatic disease pose devastating neurodevelopmental consequences for children.^16^ Importantly, chemotherapy is not always effective, and many compounds that exhibit promising activity *in vitro* fail to treat patient tumors *in situ*. The BBB and BSCB are a major obstacle to effective therapy for brain tumor patients, as many therapeutic agents are not able to reach the CNS parenchyma or require unacceptably high doses that pose toxicity to peripheral organs.

Nanoparticle encapsulation offers advantages for drug delivery through sustained release and redirection of drug fate within the body, which can improve tolerability, reduce toxicity, and enhance tissue-specific bioavailability of therapeutic agents.^17^ Although nanoparticle systems have been a central focus for therapeutic approaches aimed at treating CNS disease for several decades, we and others have reasoned that the BBB remains a primary obstacle to parenchymal delivery following nanoparticle administration by the intravenous route.^18,19^ We found that large, solid polymer nanoparticles typically remain trapped on the luminal aspect or behind the basement membrane of the BBB even when optimized with BBB-targeting ligands and even when apparent enhancements in small molecule delivery are achieved.^18^ Although delivery of large nanoparticles across the BBB is not impossible, growing knowledge in the field recognizes that much of the CNS delivery achieved by systemically circulating nanoparticles can occur via passive diffusion of lipophilic molecules, rather than by nanoparticle transit into the parenchyma; this ultimately limits the spatial distribution of drug.

Intrathecal delivery, i.e., substance administration directly into the CSF, is an attractive strategy for achieving compartmental delivery throughout the CNS.^12^ Intrathecal drug delivery, including intrathecal chemotherapy, is not a new concept;^20,21^ however, the development of polymeric nanoparticles for delivery to the CSF is less well studied, particularly at a mechanistic level. We have previously demonstrated that intrathecally administered nanoparticles distribute rapidly within the subarachnoid space to reach tissues that are distant from the injection site,^8^ and we have also examined how the route of administration (lumbar vs cisterna magna) affects the delivery of model nanoparticles to the CNS.^22^ Here, we hypothesized that encapsulation of the lipophilic HDACi Panobinostat within the CDN polymer system would improve panobinostat delivery to CNS tissues after intrathecal administration, which we predicted would facilitate treatment of patient-derived, orthotopic MB exhibiting LM in mice.

Colloidal stability is an essential design consideration for nanomedicine in the CNS,^23^ and CSF is a biologically unique milieu relative to other fluids of the body. Unlike blood, which contains a high concentration of soluble proteins, as well as platelets, lipids, and immune cells, CSF is relatively free of substances, bearing a lower concentration of ions, proteins, and cells. Although we did not specifically predict that nanoparticles would be unstable in CSF, we have observed this problem for several polymeric nanoparticle systems (data not shown), whereby aggregation is observed in aCSF when nanoparticles are otherwise stable in pure water or typical buffers such as PBS. Other investigators have also explored this issue.^23,24^ Curtis and colleagues performed detailed mechanistic studies on polystyrene nanoparticles, which revealed a specific role for divalent cations (Mg^2+^ and Ca^2+^) in generating pH-dependent aggregation of polystyrene nanoparticles. PEGylation almost entirely abrogated this instability, suggesting that ion-pairing with surface-exposed carboxyl moieties drove this process.^23^ Wang, et al., probed composition of the protein corona for polystyrene nanoparticles with and without transferrin ligand. Importantly, their methodology involved first infusing nanoparticles into the subarachnoid space and then removing samples of CSF, which enabled analyses that are considered directly relevant to the *in vivo* circumstance. This is a powerful approach, and it revealed that targeting efficacy of the transferrin-modified nanoparticles was lost following exposure to CSF.^24^ Both of these reports focused on surface-mediated phenomena, since polystyrene is rigid and non-degradable. Nanoparticles that degrade in biological fluids or that are formulated via self-assembly will encounter additional challenges. Self-assembly systems that are not cross-linked after fabrication will always carry the possibility of degradation or disassembly *in vivo*, and it can be difficult to predict solely based on *in vitro* experiments.^25^

We first identified instability of pCDN_4_ during early-stage PET imaging experiments in mice. Radiolabeled CDN_4_ nanoparticles failed to distribute consistently within the CSF of the subarachnoid space, instead remaining immediately adjacent to the injection site (CDN_4_ data not shown). To engineer a better CDN, we focused on altering the structure of the core of the nanoparticle to generate a more compact and less mobile polymer network. The newly designed CDN_5_ network not only exhibits increased stability in *in vitro* assays but also demonstrates extensive movement within the subarachnoid space to reach distant targets, which serves as confirmation that a stable nanoparticle system is able to reach tissue regions that the original version could not. These results highlight that drug design for targeting the CNS or subarachnoid space should directly address the unique fluids and barriers of the CNS microenvironment.

Panobinostat is a promising drug to treat tumors in the CNS, which has led to its clinical testing in GBM (NCT05324501, NCT00859222, NCT00848523), DIPG (NCT04341311, NCT02717455, NCT03632317, NCT03566199, NCT05009992), and, by us, in pediatric MB (NCT04315064. Our preclinical work suggests that although Panobinostat is BBB-permeant, it is degraded rapidly by esterases (similar to other HDACi), and the bioavailable free fraction within the parenchyma is too low for the drug to be effective as a monotherapy when the drug is dosed intravenously.^6,7^ These challenges form the primary motivation for developing drug carriers that can solubilize and distribute Panobinostat more effectively to treat CNS malignancies.

Direct administration of Panobinostat to fluid compartments is severely limited by its very poor aqueous solubility. There are two clinical-grade formulations that have been used for Panobinostat. The first was the orally bioavailable tablet, Farydak™, which contains excipients that are incompatible with intrathecal administration; this formulation was also recently withdrawn from the market voluntarily due to a failure to conduct post-approval trials.^26,27^ The second formulation is MTX110, which is a cyclodextrin-solubilized version of Panobinostat that has been tested by us and others for administration to pediatric tumors via the intrathecal or intratumoral routes (NCT03566199, NCT04315064). In preclinical models, Panobinostat is often formulated for intraperitoneal injection with DMSO, highly concentrated PEG, or Tweens, which are agents that would not be considered compatible for delivery via the intrathecal route. We have previously examined the CNS distribution of Panobinostat and other HDACi by solubilizing Panobinostat in Tween and PEG, which enables intravenous administration by lateral tail vein injection in mice.^6,7,28^ These studies revealed the whole-brain brain concentration of Panobinostat peaked at ∼570nM (1 hour after treatment) and declined to ∼230nM 2 hours after treatment. ^7^ We have also studied compartmental delivery of Panobinostat to the CSF following 4^th^ ventricle administration of MTX-110 in rhesus macaques.^13^ In these studies, Panobinostat was detected in CSF sampled from the 4^th^ ventricle, peaking at ∼12µM (15 min after treatment) and declining to 0.2µM 2 hrs after treatment. In contrast, when CSF was sampled from the lumbar region, drug concentrations were much lower, peaking at ∼90nM (1 hr after treatment) and declining to ∼7nM by 2 hrs after treatment.^13^ The nonhuman primate studies establish the feasibility and safety of the intrathecal approach, especially for tumors that are located within the posterior fossa, but also suggest that the spatial distribution of this compound could be further optimized. By comparison, in the work reported here, we measured concentrations of ∼1.5µM and ∼6µM in whole brain for cyclodextrin-solubilized versus nanoparticle encapsulated Panobinostat, respectively, 2 hrs after treatment; levels in lumbar spinal cord exceeded 2µM 2 hrs after treatment. Thus, intrathecal administration of nanoparticle-formulated Panobinostat achieves a higher concentration in bulk tissue of the CNS in an acute setting when compared to cyclodextrin-formulated or intravenously-administered drug.

Taken in sum, our results present an HDACi nanoparticle formulation optimized for CNS drug delivery by the intrathecal route. Nanoparticle encapsulation enhanced delivery of both IR780 and Panobinostat to the CNS compared to freely- or cyclodextrin-solubilized forms. Administration of pCDN_5_ to mice bearing orthotopic, patient derived MB activated pharmacodynamic targets to slow tumor growth, which ultimately yielded improved survival and a lower overall incidence of bioluminescent-detectable LM prior to endpoint. These studies identify the IT-CM route of administration as a promising method to achieve effective and tolerable compartmental drug delivery to the CNS.

## 4. Methods

### 4.1 Nanoparticle formulation and characterization

#### 4.1.1 Synthesis of CDNs

Cyclodextrin network polymers (CDN-4 and CDN-5) were synthesized based on our previously described methods.^11^ Acrylated beta-cyclodextrin (acr-CD) was produced by dissolving beta-cyclodextrin (CD, 2 mmol) in N-Methyl-2-pyrrolidone (NMP) and reacting with acrolyl chloride (24.7mmol). This solution was first mixed under ice-cold conditions for 1 hr, after which it was heated to 21°C and allowed to react for 48 hours. The resultant acr-CD was precipitated in distilled water, collected, washed by filtration, and lyophilized for 48 hrs. To generate the CDN-4 polymer, acr-CD (0.015 mmol) was mixed with 1,4-butenadiol diacrylate (0.675 mmol) and amine-terminated methoxy poly(ethylene glycol) (mPEG550-amine, 0.475 mmol,) in a mixture of EtOAc and CHCl3 (3:7 ratio). To generate the CDN-5 polymer, acr-CD (0.03 mmol) was mixed with 1,6-hexanediol diacrylate (0.65 mmol) and bi-functional poly(ethylene glycol) (COOH-PEG-amine, 0.95 mmol). These solutions were heated to 65°C, allowed to react while stirring at 600 rpm for 12 hrs, and dried under vacuum. The resultant crude was dispersed into a large volume of distilled water (50mL), passed through a 0.45μm bottle-top filter to remove large aggregates, and thoroughly washed via ultrafiltration (Amicon Ultra-15 centrifugal filers, 10 kDa MWCO, 15mL capacity) for two 20 min spins at 5000 RCF. The concentrated polymer solutions were aliquoted, lyophilized, and stored at -80°C until use.

#### 4.1.2 Radiochemistry

NODA-GA grafted CDN polymers were obtained via a modified CDN synthetic protocol, employing amino-NODA-GA chelator (0.5wt% of amino-PEG_3_-COOH) in the Michael addition reaction. NODA-GA grafted polymers were formed into nanoparticles through the addition of hydrophobic payload, as described above. After self-assembly, nanoparticles were added to sodium acetate buffer (pH=6.5) and treated with radioactive ^64^CuCl_2_ for chelation. This mixture was incubated for 1 hr at 30 °C, after which radiochemical purity was assessed via thin layer chromatography (TLC) analysis of the reaction mixture with scintillation counting. The reaction mixture was further concentrated via ultrafiltration to enrich the radiochemical purity as needed for *in vivo* experiments. Colloidal stability and ^64^Cu retention studies of nanoparticles were carried out in PBS or artificial cerebrospinal fluid (aCSF) with or without 10% FBS (diluted in water).

#### 4.1.3 Nanoparticle formulation by self-assembly

Nanoparticles composed of CD, CDN-4, or CDN-5 were formulated by doping hydrophobic payload into an aqueous dispersion of polymers (2mL, 20 mg polymer). Payloads for this work consisted either of Panobinostat (20 mg) or a mixture of miglyol 812N (0.25mg) and IR780 iodide (0.25mg) in 50µL of DMSO. Aqueous dispersions of payload with polymer were maintained on a rocker at room temperature for 18 hrs, during which time hydrophobic payloads enter the polymer network to facilitate self-assembly. This is a formulation approach that we optimized in previous work.^11^ The resultant nanoparticles were concentrated via ultrafiltration (Amicon Ultra-15 filters, 3kDa MWCO, 0.5mL capacity) for four 20 min spins at 5000 RCF, lyophilized, and stored at -80°C until use.

##### 4.1.4 0 Nanoparticle characterization

Hydrodynamic diameter and zeta potential for each formulation were measured via dynamic light scattering (DLS, NanoBrook 90 Plus Zeta particle size analyzer, Brookhaven Instruments, Holtsville NY). For DLS measurements, nanoparticles (20mg/ml) were dispersed in either water (size) or KCl (zeta potential, 2mL of a 1mM aq. solution) and added to a clean cuvette for readings on a number-averaged basis. Drug loading was determined by dissolving lyophilized formulation aliquots in DMSO (5mg/mL) and reading absorbance (310 nm) on a Tecan plate reader. Arbitrary units were converted to mass through comparison to a carefully constructed control curve, whereby known concentrations of Panobinostat were spiked into control (non-drug containing) CDN solutions (10 µL drug into 40µL of CDN polymer in DMSO). Results are reported for a minimum of 3 separate readings for all characterization experiments.

#### 4.1.5 Nanoparticle stability

To assess stability, pCD or pCDN formulations were suspended in artificial CSF (aCSF) buffer and incubated on rocking at 37°C. The pH of aCSF solutions was adjusted for three conditions: low (pH = 4), neutral (pH = 7.2), and high (pH = 10.2). Small samples were removed at regular intervals to evaluate hydrodynamic diameter and zeta potential via DLS, as described in section 2.2.4.

### 4.1 Animal Experiments

C57/Bl6 albino mice used for distribution experiments were purchased from the Jackson Laboratory (#000058). NOD-SCID IL2Rγ null (NSG) mice used for tolerability studies and intracranial human tumor transplantation were purchased from The Jackson Laboratory (#005557). All experiments were performed in accordance with national guidelines and regulations with the approval of the animal care and use committees at the Sanford Burnham Prebys Medical Discovery Institute, the University of California San Diego (San Diego, CA, USA), and the University of Texas Health Science Center at Houston (Houston, TX)

#### 4.2.1 Percutaneous infusion of substances into the cisterna magna or lumbar regions

Intrathecal–cisterna magna (IT-CM) infusions were performed via a percutaneous method,^29^ which in our experience is both more experimentally reliable and substantially better tolerated by mice compared to surgical approaches. Substances were prepared no more than 2 hrs prior to administration and loaded into a 50µL NeuroSyringe™ (Hamilton Company, Reno, NV) fitted with a rubber stopper to target an injection depth of ∼3.5mm. We note that it is important to optimize the injection depth for individual experimenters and mouse strain; we validated the accuracy of our injection approach in preliminary studies utilize Evan’s Blue (data not shown). Mice were anesthetized with 2% isoflurane and positioned on a flat surface with inhaled anesthesia supplied by a nose cone. To enable percutaneous injections, mice were briefly removed from the nose cone and draped over a cylinder such that the neck achieved a high arch, which maximizes the surface area of the cisternal magna. The rostral-caudal location of the cisterna magna was precisely located by palpating between the bottom of the skull and top of the vertebral column with gloved fingers. The non-dominant hand was used to gently secure the head while the dominant hand rapidly inserted the needle tip into the desired rostral-caudal location and centered along the medial line. Injections were provided as a slow bolus, after which mice were placed in a recovery cage and observed until ambulatory. Infusion volume (5µL) and speed (slow bolus over ∼20-30 seconds) were consistent for all experiments. For lumbar administration, mice were similarly anesthetized and positioned on a flat surface. The iliac crest was identified along the spine of the mouse, and an insulin syringe was inserted between the L4 and L5 spinal segments for bolus infusion. The accuracy of the infusion location was confirmed by a corresponding tail flick.

#### 4.2.2 Maximum Tolerated Dose

Tolerability of pCD and pCDN formulations was assessed in non-tumor bearing NSG mice (n=3 per dosing group, females). pCD or pCDN formulations were administered via the IT-CM route with dose escalation as follows. For single-dose studies, we examined 2, 3 and 4µg for pCD vs 2, 3, 4, 5, 6, 7, and 8ug for pCDN. For multi-dose studies, we examined 0.5, 1, and 2ug for pCD vs 2, 3, 4, 5, 6, and 7ug for pCDN. Multi-dosing was conducted 2x/week for 4 weeks, at which point the studies were terminated. Mice were observed immediately following infusions and on a daily basis thereafter. Lack of tolerability was defined as the appearance of any neurological symptom or >20% weight loss in any mouse at the given dose.

#### 4.2.3 Tumor Inductions and Monitoring

Med-411FH PDX tumor cells engineered to express luciferase were stereotaxically implanted into the cerebellum of 8-10 week old NSG mice. Mice were anesthetized with 2% Isoflurane (2L/min, MWI Veterinary Supply). An incision was carefully made in the skin over the posterior skull, and a hole in the skull was created with a 16-gauge needle 2mm posterior to lambda at the midline. 1x10^5^ cells suspended in 5μL Neurocult media (StemCell Technologies #5750) were injected using a blunt-end Hamilton syringe (#88000) angled at 35° and targeting 3.5mm deep into the cerebellum. Tumor-bearing mice were imaged 1x/week using an IVIS Spectrum system under anesthesia after intraperitoneal luciferin injection (150mg/kg). All mice were monitored for humane endpoint morbidity or toxicity (loss of balance and coordination, >20% weight loss, inactivity), whereupon they were euthanized. To assess incidence of metastasis, we scaled all IVIS images to the same level and examined the spinal cord ROI for detectable signal. The lower limit for scaling was selected as 3e7 (radiant efficiency, photons/sec per μW/cm^2^), as this was a value that captured the dynamic range of signal across multiple study cohorts. Results are reported on a weekly basis (weeks 5, 6, and 7) as the fraction of mice still living with IVIS-detectable metastasis.

#### 4.2.4 Pharmacodynamic Studies

Cerebellum and spinal cord tumors from acutely CDN-treated Med-411FH tumor-bearing mice were dissected and enzymatically dissociated into a single cell suspension. For Western Blot, whole cell protein was purified from cerebellum tumor cells using RIPA lysis buffer (Millipore, #20-188) with protease/phosphatase inhibitors (Cell Signaling Technologies #5872). Total protein was resolved using standard SDS-PAGE and transferred to a 0.2μm PVDF membrane (Invitrogen #LC2002). 40μg of protein per lane was probed for histone 3 lysine-9 acetylation (H3K9ac) using a 1:1000 dilution of primary antibody (Cell Signaling Technology #9649) followed by a 1:1000 dilution of HRP-conjugated anti-rabbit secondary antibody (Cell Signaling Technology #7074), and detected by chemiluminescence (Clarity Western ECL Substrate, BIO-RAD #1705060). The same membrane was re-probed for BETA-ACTIN using a 1:1000 dilution of primary antibody (Cell Signaling Technology #4970) followed by a 1:1000 dilution of HRP-conjugated anti-rabbit secondary antibody (Cell Signaling Technology #7074). Images were developed using a BIO-RAD ChemiDoc Imaging System and quantified using Image Lab software (Version 6.0.1 build 34). For Quantitative Reverse Transcriptase PCR (qRT-PCR), total RNA was extracted from cerebellum or spinal cord tumor cells using the RNeasy Plus Micro Kit (Qiagen #74034). cDNA was generated using iScript Reverse Transcription Supermix (BIO-RAD #1708840). RT-PCR was performed using iQ SYBR Green Supermix (BIO-RAD #1708880) with a CFX96 Real Time PCR System (BIO-RAD). The following primers were used: FOXO1 Forward: 5ʹ-TGATAACTGGAGTACATTTCGCC-3ʹ, FOXO1 Reverse: 5ʹ-CGGTCATAATGGGTGAGAGTCT-3ʹ, BETA-ACTIN Forward: 5ʹ-TGACGTGGACATCCGCAAAG-3ʹ, BETA-ACTIN Reverse: 5ʹ-CTGGAAGGTGGACAGCGAGG-3ʹ.

#### 4.2.5 Efficacy Studies

Med-411FH tumor-bearing mice were imaged by IVIS beginning 9 days after cell implantation, at which point they were allocated into treatment groups to balance tumor sizes for initiation of treatments. IT-CM treatments began on day 12 after cell implantation and consisted of 6μg pCDN_5_ or equivalent CDN_5_ mass. Subsequent IT-CM treatments were performed 2x/week through to humane endpoint.

#### 4.2.6 Small Animal PET/CT Imaging

Positron Emission Tomography (PET) / Computed Tomography (CT) imaging studies were performed with a Siemens Inveon PET/CT multimodal system for small animals (Siemens Medical Solutions, Knoxville, TN). Scans were performed in Inveon Aquisition Workplace with the following workflow: 1) PET emission acquisition, 2) CT acquisition, and 3) PET reconstruction. CT acquisition data were collected with the following parameters: x-ray maximum voltage of 80 kV, maximum anode current of 500 μA, exposure time of 260 msec for each of 120 rotation steps over a total rotation of 220°C. PET/CT reconstructions were conducted with a 2D Ordered Subset Expectation Maximization (OSEM2D) and a Feldkamp cone-beam algorithm, using a Shepp-Logan filter. PET and CT image fusion and image analysis were performed using ASIPro, Inveon Research Workplace (Siemens Preclinical Solutions) and AMIRA (version 3.1).

#### 4.2.7 Small Animal NIRF Imaging

Near Infrared Imaging (NIRF) images were taken in live mice following PET/CT scans and fluorescence was measured along the ventral and dorsal side of the mouse. Mice were imaged at approximately 30min, 1hr, and 2hrs following injection. Relevant organs (i.e., cervical lymph nodes (CLNs), inguinal LNs, renal LNs, kidneys and spleen) were then extracted and imaged *ex vivo*. All images were acquired under identical conditions with an exposure time of 200ms.

#### 4.2.8 Preparation of tissues for MALDI

Healthy C57 mice received an infusion of 8ug pCDN_5_ or 2ug cyclodextrin solubilized Panobinostat (pCD). These treatments were provided at their respective single-dose MTD. Two hours after treatment, subjects were euthanized and the brains and spinal cords were extracted. Tissues were snap-frozen in liquid nitrogen, mounted onto a specimen chuck, and sectioned coronally at 10 µm thickness (Microm HM550, Thermo Scientific, Waltham, MA). The tissue sections were thaw-mounted onto indium tin oxide (ITO) slides. A tissue microarray (TMA) mold was used to prepare a collagen mimetic model for Panobinostat quantitation by MALDI MSI. Panobinostat concentrations ranging from 0.1 - 50 µM were spiked into rat tail collagen and pipetted into 1.5 mm core diameter channels of the 40% gelatin TMA mold and frozen at -80°C. The collagen mimetic mold was sectioned, and thaw-mounted on top of control mouse brain homogenate, adjacent to the mouse brain tissue sections for analysis. Twelve serial sections moving medial to lateral were taken every 150 μm and thaw-mounted to Bruker Big Slides (Bruker Scientific LLC, USA Part No. 8259387), with serial sections preserved for hematoxylin and eosin (H&E) staining. The ITO slide mounted with the tissue and mimetic mold sections were placed in a vacuum desiccator before matrix deposition. 2,5-dihydroxybenzoic acid (160 mg/mL) matrix was dissolved in 70:30 methanol: 0.1% TFA with 1 % DMSO and applied using a TM sprayer (HTX Technologies, Chapel Hill, NC) at a flow rate (0.18 mL/min), spray nozzle velocity (1200 mm/min), nitrogen gas pressure (10 psi), spray nozzle temperature (75 °C), and track spacing (2 mm) for two passes. Optical microscopy images of the H&E-stained serial tissue sections were acquired using a 10x objective (Zeiss Observer Z.1, Oberkochen, Germany).

#### 4.2.9 MALDI MRM mass spectrometry imaging

A multiple reaction monitoring (MRM) approach was used for quantitative imaging of Panobinostat by monitoring the transition of the precursor ion fragment to product ion (350.18°158.09) using a dual source, timsTOF fleX mass spectrometer (Bruker Scientific LLC, Billerica, MA) in positive ion mode and with acquisition between m/z 100-650. Tandem MS parameters were set for a collision energy of 23 eV with an isolation width of 3 m/z. MALDI MS images were acquired with a laser repetition set to 5,000 Hz with 2,000 laser shots per 100 µm pixel. SCiLS Lab software (version 2024a premium, Bruker Scientific LLC, Billerica, MA) was used for data visualization without data normalization. The average ion intensity for each spiked TMA sectioned core area was plotted against corresponding Panobinostat concentration from 0.1- 50 µM for calibration of the MALDI MS signal, resulting in a limit of detection (LOD) of 0.2 µM (S/N ratio of > 3), and a limit of quantification (LOQ) of 0.7 µM (S/N ratio of > 10).

## Acknowledgements

The authors gratefully acknowledge funding from the National Institutes of Health (R01NS116657 (to RWS), R01HD099543 (to RWS, RJW, ES, and NRA), R01NS111292 (to RWS and RJW), R21NS107985 (to RWS), Ian’s Friends Foundation (to RWS), and Rice University (to OB).

