## Supplemental Data for "Nanoparticle encapsulation enhances spatial distribution of Panobinostat to treat metastatic medulloblastoma via the intrathecal route"

### Supplementary Materials

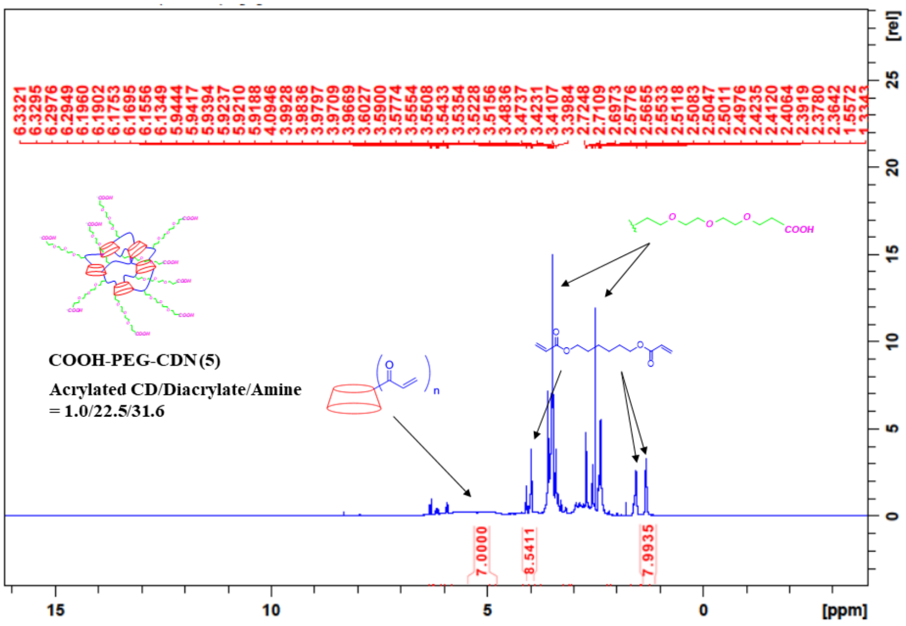


**Figure SI1**. 1H-NMR spectra of CDN-5 in DMSO-d6 recorded in Bruker Avance 500 MHz NMR spectrometer. Stoichiometric ratios of constituent units are recorded. Assignments of 1H-NMR peaks were attributed to the protons present in the individual molecular entity, which indicates to the composition of the material (CDN-5).

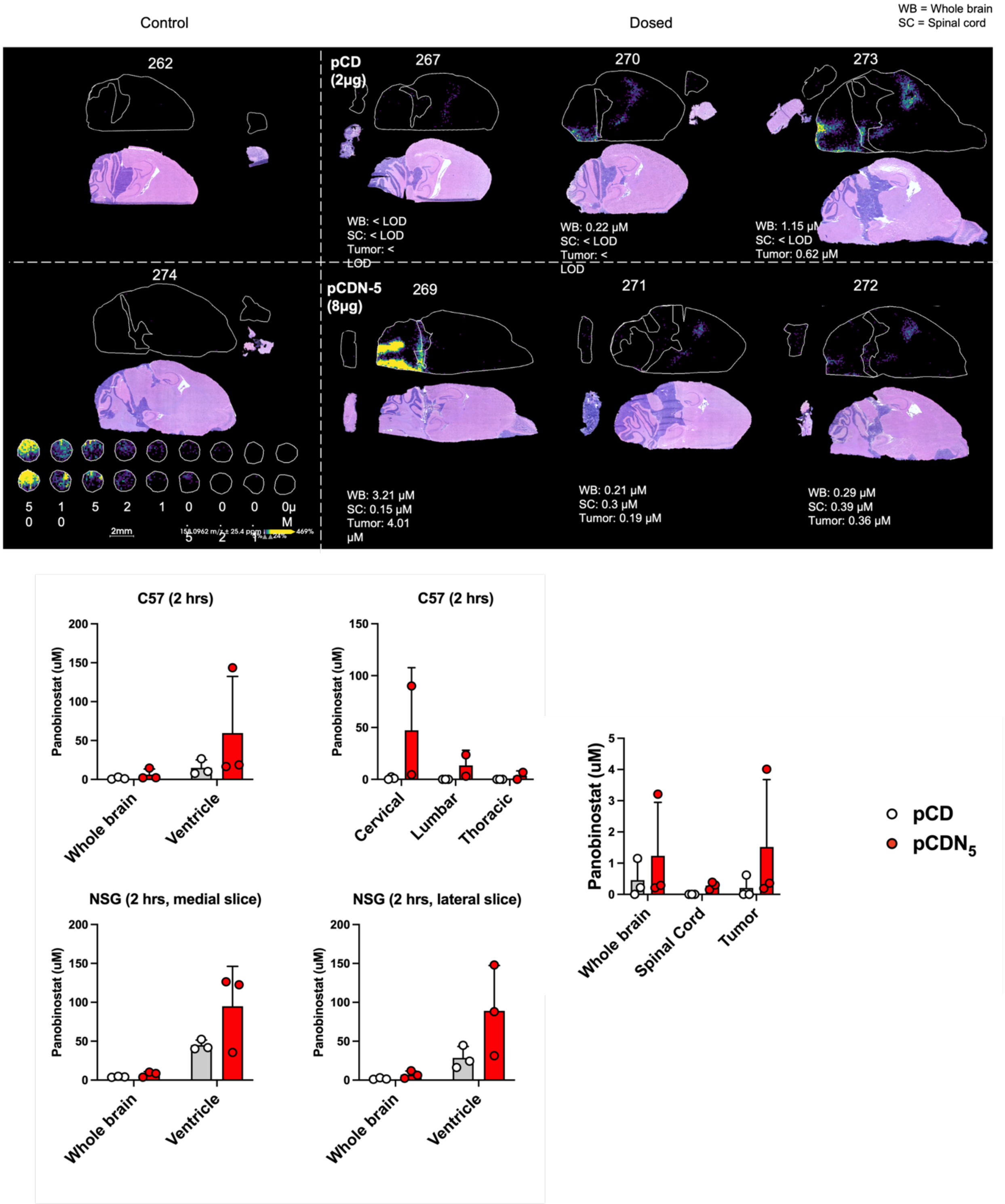


**Figure SI2**. Complete MALDI data set showing quantification of Panobinostat levels in healthy C57 and NSG mice, as well as in tumor-bearing NSG mice. All data were acquired as described in Methods.
